# AKAP8 interacting with DDX5 to regulate R-loop balance involves in lung carcinoma cell growth

**DOI:** 10.1101/2023.11.28.568985

**Authors:** Xu Wang, Liang Liu

**Affiliations:** Guangzhou National Laboratory, Guangzhou, Guangdong, 510005, China

**Author notes:** **Corresponding author:** Xu Wang, Liang Liu.

**Keywords:** lung carcinoma, AKAP8, R-loop

## Abstract

Triple-nucleotide chain of RNA:DNA hybrid structure R-loop is formed during transcription process which is closely associated with transcriptional regulation, transcriptional termination, epigenetic modifications, and structure of chromatin. Dysregulation of R-loop formation or resolution may disturb normal DNA replication or RNA transcription, leading to associated progress of disease. Herein, we found A-kinase-anchoring protein 8 (AKAP8) bind with R-loop structure in cell. Knock down of AKAP8 showed perturbation of balance of genomic R-loop formation and gene transcription. Evidences was shown that AKAP8 interacted with R-loop resolution protein ATP-dependent DEAD box RNA helicase DDX5 and increased chromatin associated DDX5 level. Finally, AKAP8 promoted UCP2 transcription and resolved R-loop level of its promoter region, and contributed to cell growth of adenocarcinomic human alveolar basal epithelial cell line A549, which may provide clues for poor prognosis of lung adenocarcinoma in high AKAP8 expression patients.

## Introduction

The R-loop is composed of tri-strand nucleic acid structure formed during transcription process. One chain of the double strand DNA is replaced by RNA chain complementary to the other strand DNA chain to form an RNA:DNA hybrid along with a exposed single DNA strand. It is thermodynamically more stable than double strand DNA (Thomas et al. 1976). R-loop is well knowledge as double edged sword for its advantages and disadvantages for cell metabolism. On the one hand, R-loop plays roles in regulation of transcription by multiple mechanism, like participate in promoter methylation, transcription factor binding, or transcription termination (Grunseich et al. 2018) (Chen et al. 2015) (Boque-Sastre et al. 2015). In addition, R-loop is indispensable for replication of bacterial plasmid and human mitochondrial genome (Renaudin et al. 2021). Also, it is necessary for process of animal immunoglobulin gene class switch recombination to generate types of antibody (Refaat et al. 2023).

On the other hand, the disadvantage of R-loop mainly addressed by its destruction to DNA. R-loop makes DNA more sensitive to damages, including transcription dependent DNA recombination, double strand DNA breaks, fragile site instability or even severe chromosome loss (Groh and Gromak 2014). Since R-loop regulates transcription profile of cell, unexpected dysregulation of R-loop by abnormal accumulation or resolution is closely associated with disease in some cases. Mutation of RNA:DNA helicase SETX cause neurodegeneration in adolescent predominant amyotrophic lateral sclerosis type 4 (ALS4) (Kannan et al. 2022). Cell of autoimmune disease Aicardi-Goutieres syndrome (AGS) accumulates R-loop and significantly activates cGAS-STING pathway, which is regarded to associate with RNase H mutation responding for cellular resolution of RNA:DNA (Lim et al. 2015) (Crow and Manel 2015). Mutation driving metabolism perturbation and R-loop accumulation induced DNA damage or blockade of transcriptional complex by breast cancer susceptibility factor BRCA is sort of related to breast tumorigenesis (Tan et al. 2017). In immune cells, unexpected R-loop accumulation simultaneously in immunoglobulin switch gene and its translocation partner including oncogene c-MYC enhance the pathological translocation of them performed by activation induced cytosine deaminase (AID) (Ruiz et al. 2011). Therefore, identification and revelation of R-loop associated protein can provide a basis for elucidation of R-loop regulatory mechanism and diagnostic strategy for disease.

Helicases are kind of family involving in R-loop metabolism. So far, several important helicases including RNA/DNA helicase Senataxin SETX, Auarius AQR, DNA helicase RECO5, RNA helicase DDX1, DDX5, DDX19, DDX21, DHX9 participate in R-loop formation and resolution (Ribeiro de Almeida et al. 2018) (Mersaoui et al. 2019) (Chakraborty et al. 2018). DEAD-box (DDX) protein containing conserved motifs including eponymous DEAD motif (Asp-Glu-Ala-Asp, D-E-A-D) are defined as largest family of double strand RNA helicase (Linder and Jankowsky 2011). DDX5 as coregulator for cellular transcription and splicing, and participator in processing of small noncoding RNA is previously shown to unwind RNA:RNA, RNA:DNA, and R-loop in vitro (Xing et al. 2017) (Mersaoui et al. 2019). Deficiency of DDX5 accumulates R-loop at propensity loci to form such structure in U2OS cell line. It is reported DDX5 is associated with XRN2 exoribonuclease to resolve R-loop resulting in release of RNA

Pol II from transcriptional termination regions (Mersaoui et al. 2019). After, some co-factor of DDX5 involving in its R-loop resolution are claimed including ATPase family AAA domain-containing protein 5 (ATAD5) restricting R-loop formation at replication fork, Yamanaka factor SOX2 inhibiting R-loop resolvase activity of DDX5, thyroid hormone receptor-associated protein 3 (Thrap3), topoisomerase III-beta (TOP3B) and THO complex subunit 5 homolog (THOC5), Treacle ribosome biogenesis factor 1 (TCOF1) promoting R-loop resolution (Kim et al. 2020) (Li et al. 2020b) (Kang et al. 2021) (Saha et al. 2022) (Polenkowski et al. 2023) (Nie et al. 2023). The resolvase activity of DDX5 is also dependent on its post modification. Protein arginine methyltransferase 5 (PRMT5) methylates DDX5 at its RGG/RG motif which promotes DDX5 interaction with XRN2 and repression of cellular R-loops (Mersaoui et al. 2019). Therefore DDX5 and associated proteins cooperate to regulate R-loop formation and resolution during genetic process. The unexpectable aberrant expression of them may impair cellular metabolism and contribute to disease progression. Sox2 interacting with DDX5 inhibits the R-loop resolvase activity of DDX5 to thus facilitate reprogramming (Li et al. 2020b). In Gastric cancer (GC), aberrantly high expression of TCOF1 cooperating with DDX5 contribute to maintaining GC cell proliferation though alleviating R-loop associated DNA replication stress (Nie et al. 2023). Besides of its function in R-loop, DDX5 resolves G4 structure of oncogene MYC promoter to regulate its transcriptional activation (Wu et al. 2019). Positive prognostic factor DDX5 expression is negatively associated with MYC expression in lung cancer (Cui et al. 2021).

In this study, we identified R-loop binding protein A-kinase anchoring protein 8 (AKAP8, also known as AKAP95) was associated with DDX5. AKAP8 is member of AKAP family protein with three classical domain of protein kinase A (PKA) binding domain exerting by hydrophobic face of conserved amphipathic helix, targeting sequence serving to tether the complex to specific subcellular compartment, and signal molecular binding domain elevating second messenger cyclic AMP (cAMP) (Langeberg and Scott 2005). Most of AKAPs reside in the cytoplasm except several members including AKAP8, AKAP8 homologue AKAP8L, and splicing factor SFRS17A (known as AKAP17A) (Kubota et al. 2015) (Martins et al. 2003) (Jarnaess et al. 2009) . For AKAP8, it is reported it regulates nuclear PKA though local cAMP from organized nuclear microdomain consisting of AKAP8, PKA, PDE4D5 (Clister et al. 2019). Besides of its classical kinase anchoring function, AKAP8 capacity of binding with RNA involves in transcription and splicing regulation. AKAP8 and selected partner hnRNPs cooperated to regulate alternative splicing (Hu et al. 2016) (Li et al. 2020a). In vitro and in cell, AKAP8 formed liquid like condensate. The regulatory ability of AKAP8 in transcription and splicing requires its special biophysical property of formed condensate with properly dynamic liquid-liquid phase separation and exchange (Li et al. 2020a). AKAP8 depleted cells shows accelerated EMT and enhanced breast cancer metastatic potential (Hu et al. 2020). The amplification and upregulation of AKAP8 in triple negative breast cancer (TNBC) are found to be correlated with poorer patient survival and its deficiency reduced growth of TNBC cell line (Li et al. 2020a). AKAP8 is determined to interact with connexin 43 (Cx43) in a dynamic pattern during cell cycle progression in lung carcinoma cell line by which regulated cell proliferation (Chen et al. 2016) (Chen et al. 2020). However, the pathophysiologic roles of AKAP8 is so far poorly undertaken.

Herein, we identified AKAP8 was R-loop association protein. Further deciphering interaction between AKAP8 and DDX5, we claimed that AKAP8 slightly enhance chromatin associated DDX5 level and regulated R-loop resolution resulting in transcription changes. It was also suggested this regulation pattern of AKAP8 may contribute to lung carcinoma cell growth offering it as potential prognostic target for lung cancer.

## Results

### HBD-TurboID for identification of R-loop associated protein

HBD is responding for RNA:DNA hybrid recognize from RNase H N terminal domain. TurboID were recombined expression with HBD and nuclear location sequence to construction for proximity biotinylation of RNA:DNA hybrid associated protein in cell nucleus (Figure 1A). HBD-TurboID were stable expressed in 293T (Figure 1B). After biotin incubation, the protein of HBD-TurboID 293T cell line showed significant biotinylation comparable to that of TurboID 293T cell line detected by Sa-HPR (Figure 1C). The biotinylated proteins were affinity precipitated (AP) by Sa beads and estimated by Sa-HPR. After incubation with biotin, more affinity precipitated proteins were detected for HBT-TurboID 293T cell line as well as TurboID 293T cell line comparing with that of without biotin (Figure 1D). The results showed functional biotin labeling for HBT-TurboID system.

**Figure 1.**
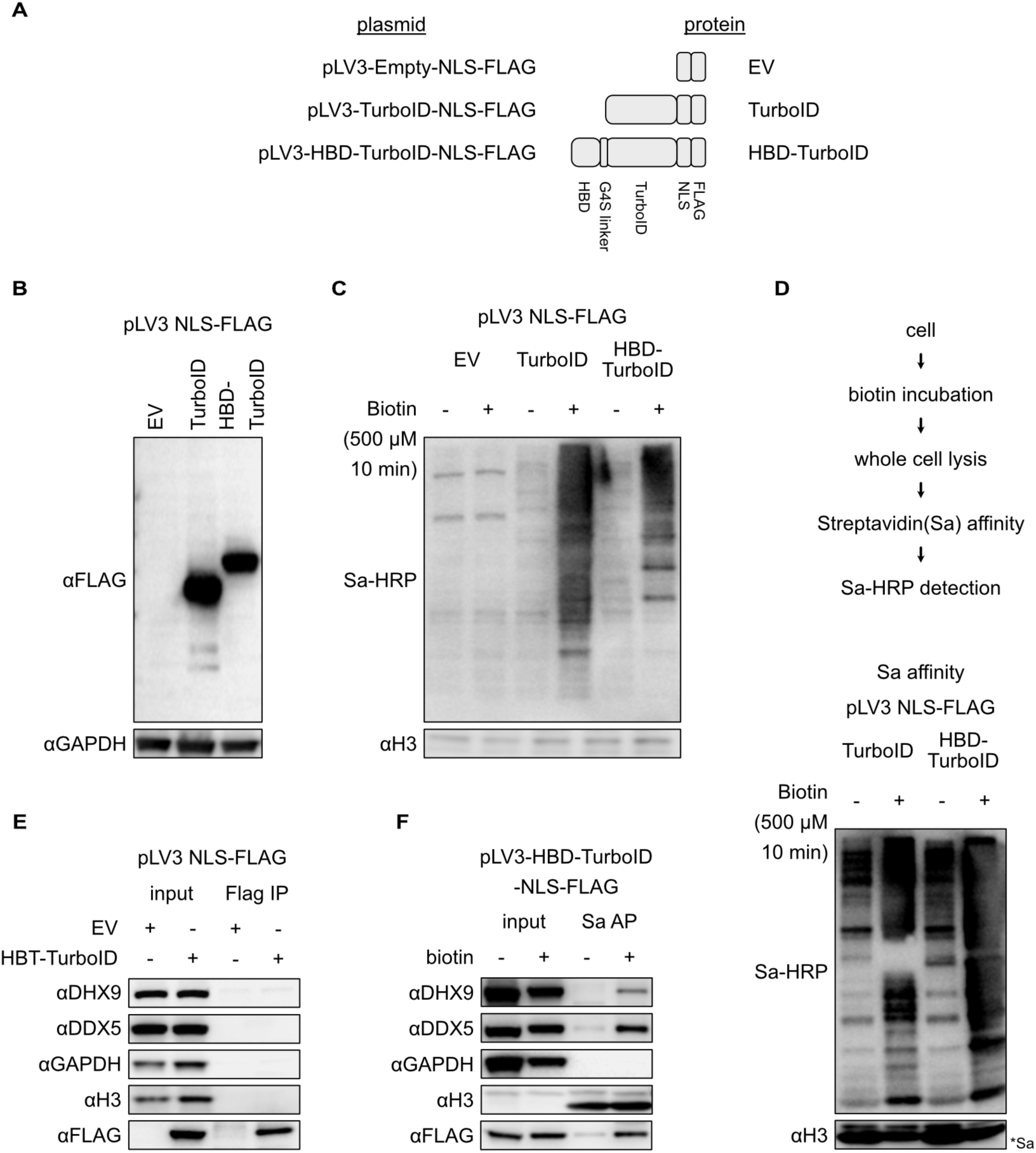
Proximity labelling of R-loop associated protein by HBD-TurboID. A, Schematic graph represented regions of HBD-TurboID protein. HBD and TurboID were recombined by G_4_S linker. NLS and FLAG were recombined in N terminal. B, HBD-TurboID protein expression in 293T stable cell line. C, Estimation of biotinylated whole cell protein by TurboID. D, Sa affinity precipitation (Sa AP) of biotinylated proteins labeled by TurboID system. Sa AP product of biotin linked protein were estimated by PAGE and Sa-HRP. E, Estimation of HBD-TurboID interaction with R-loop associated proteins and reference protein by FLAG IP. F, Estimation of R-loop associated protein by HBD-TurboID system.

HBD-TurboID were designed to explore R-loop associated proteins. To exclude if direct interaction of HBD-TurboID and those proteins exist, HBD-TurboID immunoprecipitation (IP) were firstly carried out and representative R-loop protein were estimated. The result showed DHX9 DDX5 as well as subcellular markers GAPDH and histone H3 were found no immunoprecipitation with HBD-TurboID protein (Figure 1E). Comparing with no biotin treatment, significantly amount of DHX9 and DDX5 instead of subcellular marker GAPDH and histone H3 were affinity precipitated by Sa beads after biotin labeling for HBD-TurboID 293T cell line (Figure 1F). HBD-TurboID system was suggested to be a efficient R-loop associated protein labeling system and its advantage was discussed in last section.

### AKAP8 binding and remodulating R-loop

AKAP8 binds with both DNA and RNA. The interactome of AKAP8 contains R-loop associated proteins including RPA, hnRNPs, and DDX (Figure S1A). To investigate the relationship between AKAP8 and R-loop, we first confirmed their interaction by HBD-TurboID proximity labelling system. Extraction of HBD-TurboID 293T stable cell line incubated with biotin were affinity precipitated by Sa beads and AKAP8 was estimated in affinity precipitated product. The result showed AKAP8 was not detected in IP product of HBD-TurboID expressed cells, however was obviously detected in AP product of HBD-TurboID expressed cell after incubation with biotin (Figure 2AB). The interaction between R-loop and AKAP8 was further confirmed by S9.6 immunoprecipitation. As is shown, AKAP8 co-precipitated with S9.6 in 293T cell while homologous protein AKAP8L did not (Figure S1B). After RNase H treatment, no AKAP8 signal was showed in S9.6 IP products of 293T cell (Figure 2C). The results suggested that AKAP8 is a new revealed R-loop associated protein. We further estimated and confirmed this interaction in A549, HeLa, U2OS, and UMSC (Figure 2DF). Giving AKAP8 was reported as RNA binding protein, we also investigated whether RNA is in charge with the interaction between AKAP8 and R-loop containing RNA. Interestingly, after RNase A treatment, more AKAP8 signal seamed shown in S9.6 IP products of various cells (Figure 2DF). This may be cause by release of AKAP8 trapped by RNA and more amount of AKAP8 interaction with R-loop.

**Figure 2.**
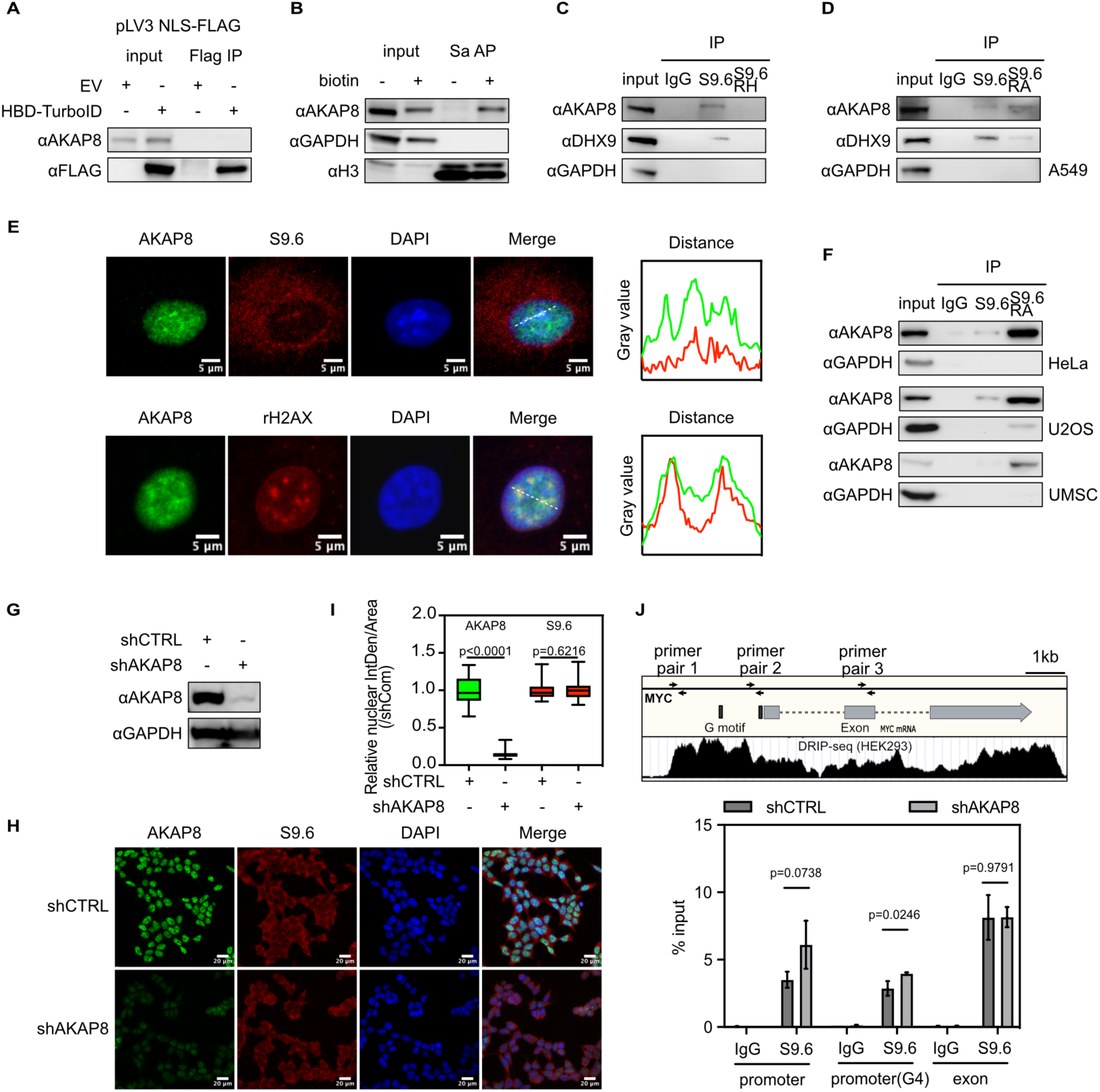
AKAP8 bind with R-loop and regulated its formation. A, Estimation of HBD-TurboID interacting with AKAP8 by FLAG IP. B, AKAP8 interacting with R-loop by HBD-TurboID system. C, AKAP8 interacting with R-loop by S9.6 IP. RH: for RNase H treatment. D, F, investigation of AKAP8 interacting with R-loop in A549, HeLa, U2OS, UMSC cells by S9.6 IP. RA: for RNase A treatment. E, immunofluorescence and signal profile of AKAP8, R-loop, rH2A.x in cells. G, Estimation of AKAP8 expressed in shCTRL and shAKAP8. H, I, immunofluorescence and signal profile of AKAP8 and R-loop (S9.6). J, diagram for MYC gene, mRNA and R-loop signal from database and evaluated by DRIP-qPCR. Primer paire 1 2 3 used for promoter, promoter (G4) and exon qPCR analysis.

Subcellular location of AKAP8 and R-loop was subsequently to be estimated. The immunofluorescence image showed AKAP8 had mostly nuclear location, while R-loop was also found partially nuclear location. Signal abundance distribution presented by gray value-distance showed accordingly (Figure 2E). S9.6 was reported to positively closed related to double strand break DNA represented by molecular marker rH2A.X. We also estimated subcellular distribution of AKAP8 and rH2A.X. It was imaged that AKAP8 formed speckles like rH2A.X foci. Interestingly, they showed obviously merged signal distribution according to gray value-distance analysis (Figure 2E). These results implicated nucleus colocation of AKAP8 and R-loop.

We nest explored whether AKAP8 involved in R-loop homeostasis. Relative nuclear average inter density of S9.6 was estimated in control and AKAP8 knock down 293T cell lines. The result showed AKAP8 signal of its knock down cell was significantly decreased in according with expression level, however bare differences for nuclear S9.6 signal comparing to control cell (Figure 2 GHI). R-loop level of MYC gene loci were investigated. The results showed that promoter(G4) region instead of exon loci of MYC gene accumulated R-loop (Figure 2J). These results suggested that AKAP8 was closely associated with R-loop in nuclear and contributed to its resolution in some region instead of general genome.

### AKAP8 interacting with DDX5 by N terminal region containing nucleus location sequence

Previous study revealed AKAP8 interactome contains R-loop associated proteins. DDX5 was one of these proteins directly participating in R-loop resolution by its ATP dependent RNA helicase activity. To investigation association of AKAP8 and DDX5, GST pull down experiment were carried out and results showed exogeneous expressed AKAP8 co-precipitated with endogenous DDX5 both in 293T and A549 cell (Figure 3A). The estimation of AKAP8 and DDX5 interaction by exogenous expression, half endogenous expression of HA-AKAP8 DDX5-FLAG confirmed interaction of these two protein in both experiments (Figure S2AB). In addition, their interaction was found to be DNA/RNA independent implicated by treatment of benzonase nuclease for cell lysate before IP process (Figure 3B Figure S2C). Then AKAP8 and DDX5 were found to be co-precipitated in 293T cell (Figure 3C Figure S2D). GST-AKAP8 and DDX5-His were also expressed in E.coli and co-precipitation was estimated by GST pull down. DDX5-His was found to be co-precipitated with GST-AKAP8 instead of GST (Figure 3D). These results revealed the direct interaction of AKAP8 and DDX5 protein.

**Figure 3.**
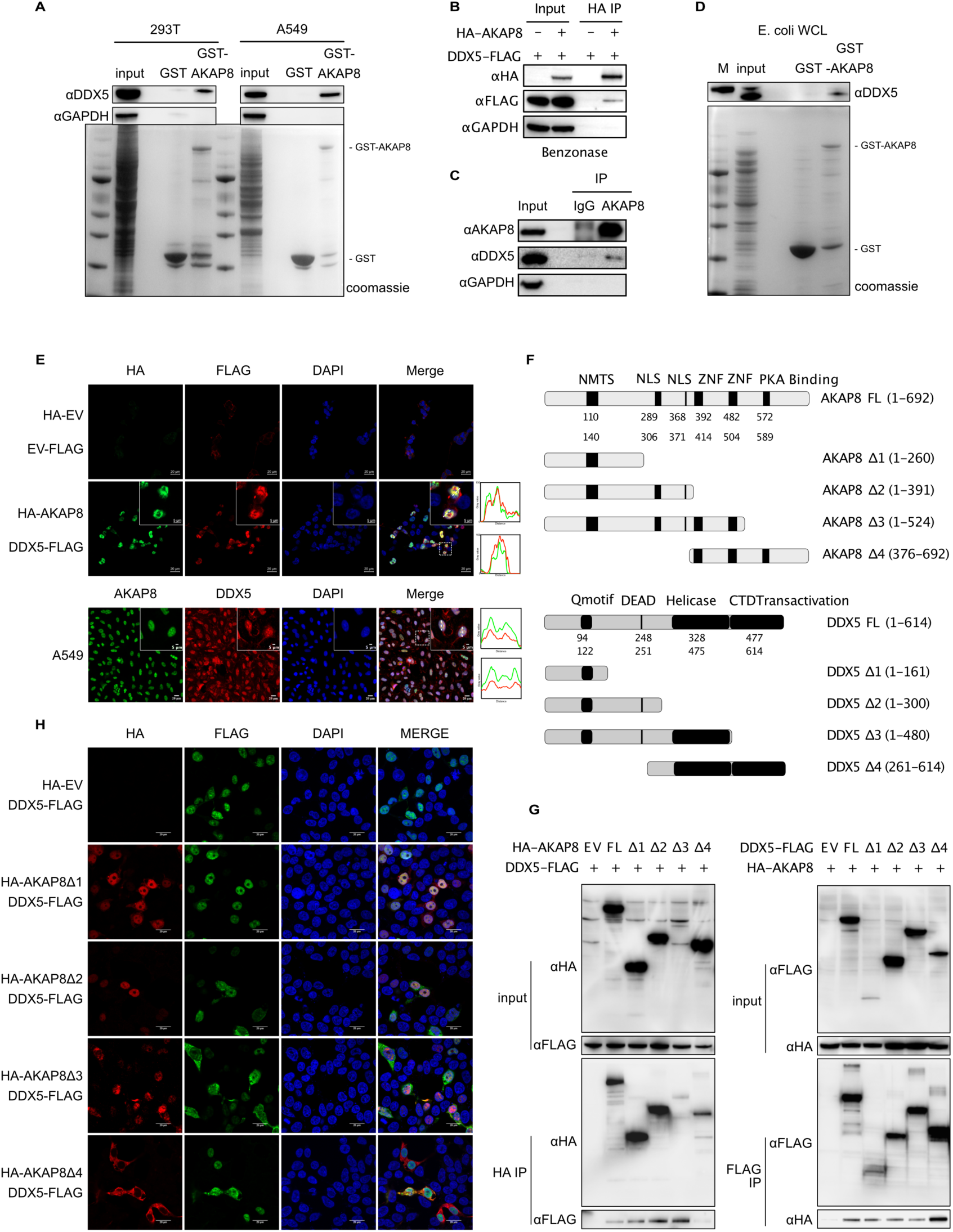
AKAP8 bind with DDX5 in nucleus. A, GST pull down estimation of AKAP8 interacting with DDX5 in 293T and A549. Input: whole cell extraction. B, AKAP8 interacting with DDX5 by HA IP after benzonase treatment. C, interaction of AKAP8 and DDX5 by IP. D, estimation of GST-AKAP8 and His-DDX5 interaction expressed in E. coli. Input: whole cell extraction of His-DDX5 expression E. coli after IPTG induction.E, immunofluorescence image of AKAP8 and DDX5. F, schematic diagram of structure domain and truncated regions of AKAP8 and DDX5. G, interacting region estimated by truncated HA-AKAP8 and DDX5-FLAG through IP. H, immunofluorescence image of truncated HA-AKAP8 and full length DDX5-FLAG expressed in 293T.

The subcellular location of AKAP8 and DDX5 was nest investigated. The immunofluorescence image of exogenous HA-AKAP and DDX5-FLAG of 293T cell showed that both of them had almost nucleus location. Besides, the signal profile distribution seemed to have a positive co-relation (Figure 3E). The immunofluorescence image of AKAP8 and DDX5 of A549 cell showed both of them had almost nucleus location and correlation of signal profile distribution (Figure 3). It was suggested AKAP8 and DDX5 were almost transported into nucleus and had a direct interaction.

To investigate binding domain of AKAP8 and DDX5, several truncated HA-AKAP8 and DDX5-FLAG taking their conserved domain into consideration were expressed in 293T cell. For human AKAP8 protein, it contains NMTS, two NLS, two ZNF, and PKA binding domain. For human DDX5 protein, it contains Q motif, DEAD box motif, helicase C terminal, and transactivation domain (Figure 3F). The HA IP of co-expressed truncated HA-AKAP and DDX5-FLAG gave the evidence that N terminal of AKAP8 contain NMTS was co precipitated with DDX-FLAG. However, the FLAG IP of co expressed truncated DDX5-FLAG and full length HA-AKAP8 showed all expressed truncated DDX5-FLAG co precipitated with HA-AKAP8 (Figure 3G). It was suggested NMTS domain of AKAP8 was responsible for their interaction however DDX5 had possibly multiple regions to bind with AKAP8.

Given subcellular colocation and interaction relation, we nest investigated whether truncated HA-AKAP8 disturb DDX5-FLAG subcellular location when co express in 293T. The immunofluorescence image showed N terminal of AKAP8 containing NMTS was sufficient to satisfy nuclear location nevertheless C terminal of AKAP8 lacking NMTS and NLS domain showed cytoplasmic distribution (Figure 4H). Overexpressed DDX5-FLAG with empty HA and other truncated AKAP8 plasmid in 293T cell showed mostly nuclear sublocation excepted for only PKA binding domain deleted AKAP8. This truncated AKAP8 showed nuclear location, but the DDX5-FLAG appear to have more cytoplasmic distribution (Figure 4H). It was imaged that N terminal of AKAP8 contributed to its nuclear targeting character and NMTS domain containing region was sufficient to achieve as well as AKAP8 interaction with DDX5. The disputation of PKA binding domain of AKAP8 may perturb DDX5 nuclear targeting.

**Figure 4.**
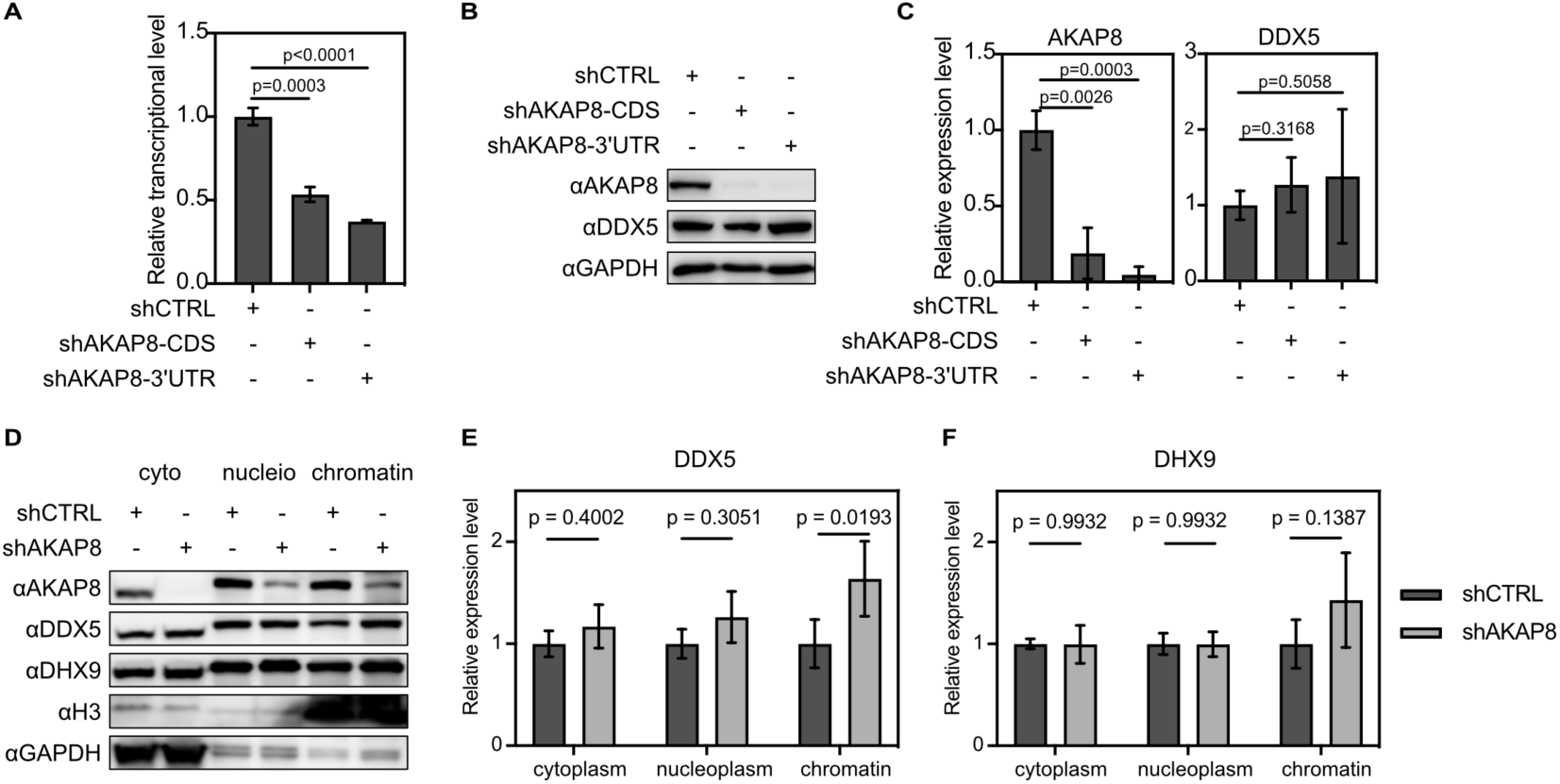
AKAP8 knock down increased chromatin DDX5 level. A, relative transcriptional level of AKAP8 in shCTRL and shAKAP8 293T cell line. B, AKAP8, DDX5 expression in shCTRL and shAKAP8 293T cell line. C, Statistic of AKAP8, DDX5 expression. D, expression of proteins in cytoplasm (cyto), nucleioplasm (nucleio) and chromatin associated (chromatin) fraction of cell. E, F, Statistic of AKAP8, DDX5 expression in cell fractions.

### AKAP8 reduction enhancing chromatin associated DDX5

To investigate whether AKAP8 affects DDX5 expression, AKAP8 knock down 293T cell lines were established. Transcriptional and expressional level of AKAP8 was significantly reduced in two cell lines (Figure 4A-C). The expression level of DDX5 was found no significant changes (Figure 4BC). Since truncated AKAP8 may perturbed DDX5 nucleus signal, we estimated subcellular DDX5 level in fractions. The results showed that DDX5 level in cytoplasm and nucleoplasm have few changes but increasing of chromatin associated component in AKAP8 depleted cell comparing to control cell (Figure 4DE). Another known R-loop protein DHX9 has bare changes in all test cell fraction (Figure 4DF). The result implicated that reduction of AKAP8 increased chromatin associated DDX5 level.

### AKAP8 playing roles in regulating oxide process of cancer cells

Given the association of AKAP8 and R-loop, and study of AKAP8 participating in splicing process, the RNA-seq by long reads method and data analysis of AKAP8 knock down cancer cell line were carried out. Comparing to control cell, differential expressed genes (DEGs) of 214 up regulated and 221 down regulated were found in shAKAP8 cell (Figure 5A Figure S3A). Tendency of relative RNA level of AKAP8 and several DEGs in two cell line were estimated by qPCR and showed accordance with reads abundance of RNA-seq (Figure 5B). Since AKAP8 was reported to play roles in RNA splice, we also investigated different expressed transcripts and alternative polyadenylation (APA). The APA motifs were analysis and listed including canonical AATAAA motif as well as CG abundance motif like CCAGCCTG (Figure S3B). The alternative splicing happened mostly by exon skipping which was over 60% of all splicing events. Other splicing ways including 5’, 3’ stie splicing ranged from about 3∼13% (Figure 5D). The different splicing ways of two cell lines showed comparable percentage value suggested AKAP8 appeared to no preference for alternative splicing location in this test condition.

**Figure 5.**
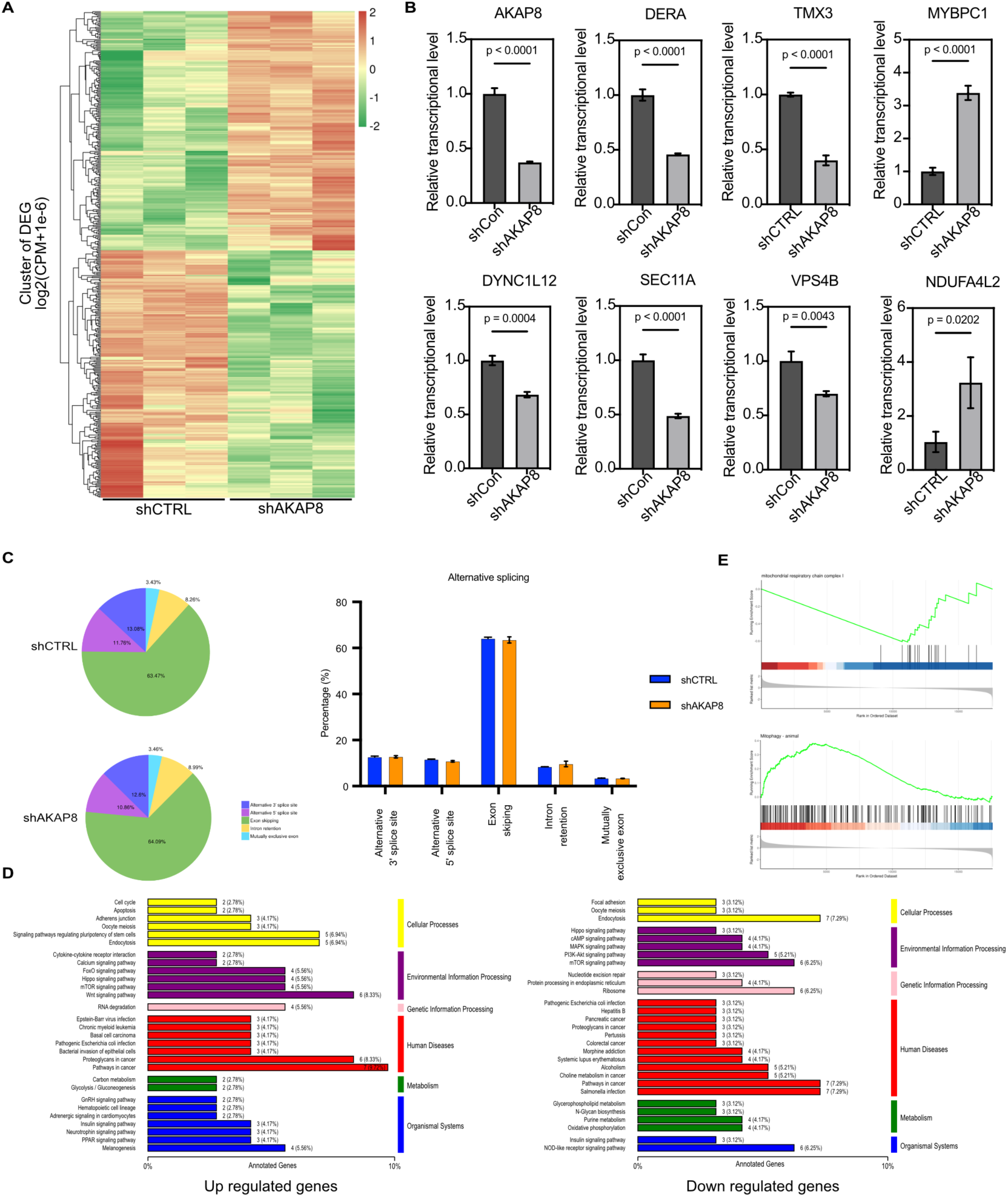
RNA-seq revealed AKAP8 regulating pathways. A, Heat map of clustered DEGs in shCTRL and shAKAP8 cell samples. B, qPCR estimation for relative transcriptional level of selected DEGs. C, statistic for alternative splicing ways for shCTRL and shAKAP8 cell. D, KEGG pathway analysis of up regulated and down regulated DEGs. F, GSEA analysis revealed AKAP8 knock down regulated mitochondrial metabolism pathways.

To comprehensively investigate roles of AKAP8 in biological metabolism and metabolic network, KEGG annotation and classification of DEGs were carried out. Classified KEGG item of upregulated and downregulated DEGs were listed respectively (Figure 5C). it was attractive that pathway in cancer item was found both in upregulated gens and downregulated genes. RNA degradation item was found only in up regulated genes while oxidative phosphorylation item was found in downregulated genes possibly contributed by transcription factor regulating pattern (Figure S3C). Gene set enrichment analysis (GSEA) of all genes were perform to achieve enrichment analysis on all genes based on the expression of all genes without prior experience . Among the GSEA item, RNA degradation item appeared to show positive enrichment score (Figure S3D). Mitochondrial respiratory chain complex I and mitophagy item related to oxidative phosphorylation pathway showed negative and positive enrichment score from comprehensive judgements respectively (Figure 5E). These GSEA character was in according with KEGG classification results. It was implicated that AKAP8 seemed to involving in RNA degradation and mitochondrial metabolism in carcinoma cell line.

### AKAP8 as a potential oncogene involving in lung adenocarcinoma progress

To investigate whether AKAP8 is associated to carcinoma progress, statistical analysis of its expression across different types of cancers were carried out. As is shown, except pancreatic adenocarcinoma (PAAD), clear cell of renal cell carcinoma (RCC), and uterine corpus endometrial cancer (UCEC), AKAP8 expression in other types of carcinoma had higher score than that in normal tissue (Figure 6A). Since over expression of AKAP8 in lung carcinoma was reported in several cases but no experimental evidences were addressed, we emphasized to investigate its roles in lung carcinoma. As is shown, both of expression of AKAP8 in a lung adenocarcinoma and lung squamous cell carcinoma tissue had significantly high score than that in normal tissue (Figure 6B). High expressed AKAP8 lung carcinoma patient seemed to have lower survival probability (Figure 6C). we investigated relation of AKAP8 and lung carcinoma cell line A549. More colony of A549 with higher AKAP8 expression formed (Figure 6D). It was implicated that high expression of AKAP8 may involve in lung carcinoma. These results suggested that AKAP8 was possibly regarded as one of oncogenes for lung carcinoma.

**Figure 6.**
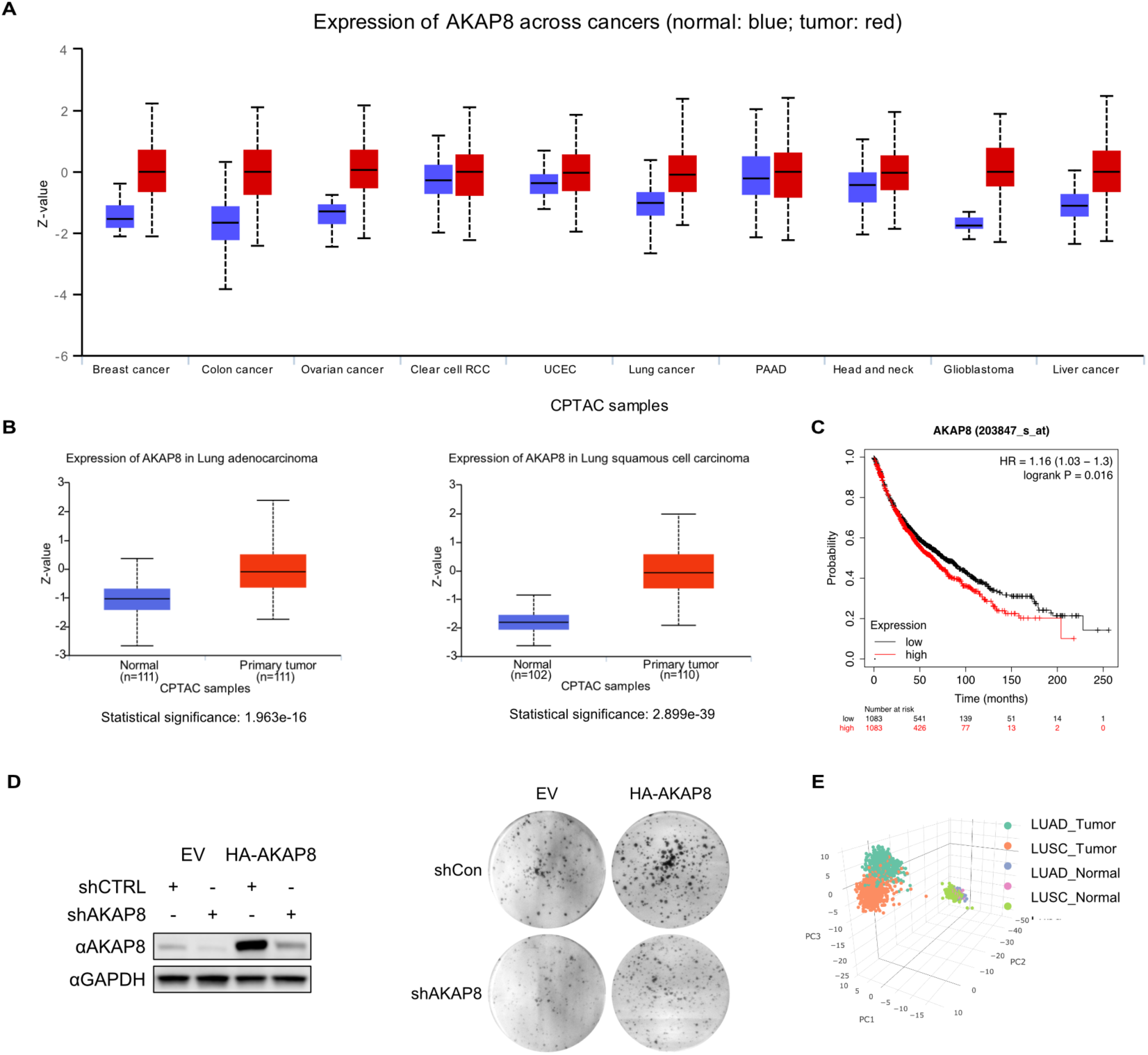
AKAP8 participated in cell growth and progress of lung carcinoma. A, Z-value estimation of AKAP8 expression across types of cancer. Blue for normal tissue; red for tumor tissue. B, Statistical analysis of AKAP8 expression in lung adenocarcinoma and lung squamous cell carcinoma by comparing that of normal tissue and tumor tissue. C, Survival probability with high and low AKAP8 expression. D, AKAP8 expression level estimation and colony formation for four cells. E, PCA statistics for DEGs set in lung carcinoma tumor and normal tissue.

We further performed PCA dimensionality reduction of DEGs of shCTRL and shAKAP8 in lung carcinoma of TCGA gene expression database. The results showed that expression profile of DEG set of normal tissue including lung, lung adenocarcinoma (LUAD) and lung squamous cell carcinoma (LUSC) (Figure 6E). Normal tissue aggregated in similar location of PCA 3d axis graph. expression profile of DEG set of LUAD and LUSC aggregated in distinct location. Moreover, the PCA aggregation of LUAD and LUSC separated in a 2d axis graph (PC1-PC3). This was implicated that AKAP8 may had difference effect of metabolism in LUAD and LUSC.

### AKAP8 remodulating R-loop accumulation and transcription of UCP2

Previous results in this study showed AKAP8 contributed to mitochondrial metabolism. One of most regulated gene of mitochondrial associated DEGs is UCP2. We found lower survival probability was presented in UCP2 high expression patients (Figure 7A). In addition, gene expression correlation analysis of AKAP8 and UCP2 was with significant p value (0.018) and positive R value. However, no significant correlation appeared to be found in both LUSC and normal tissue (Figure 7B). It is implicated that the positive regulation of UCP2 by AKAP8 may play roles in LUAD progress.

**Figure 7.**
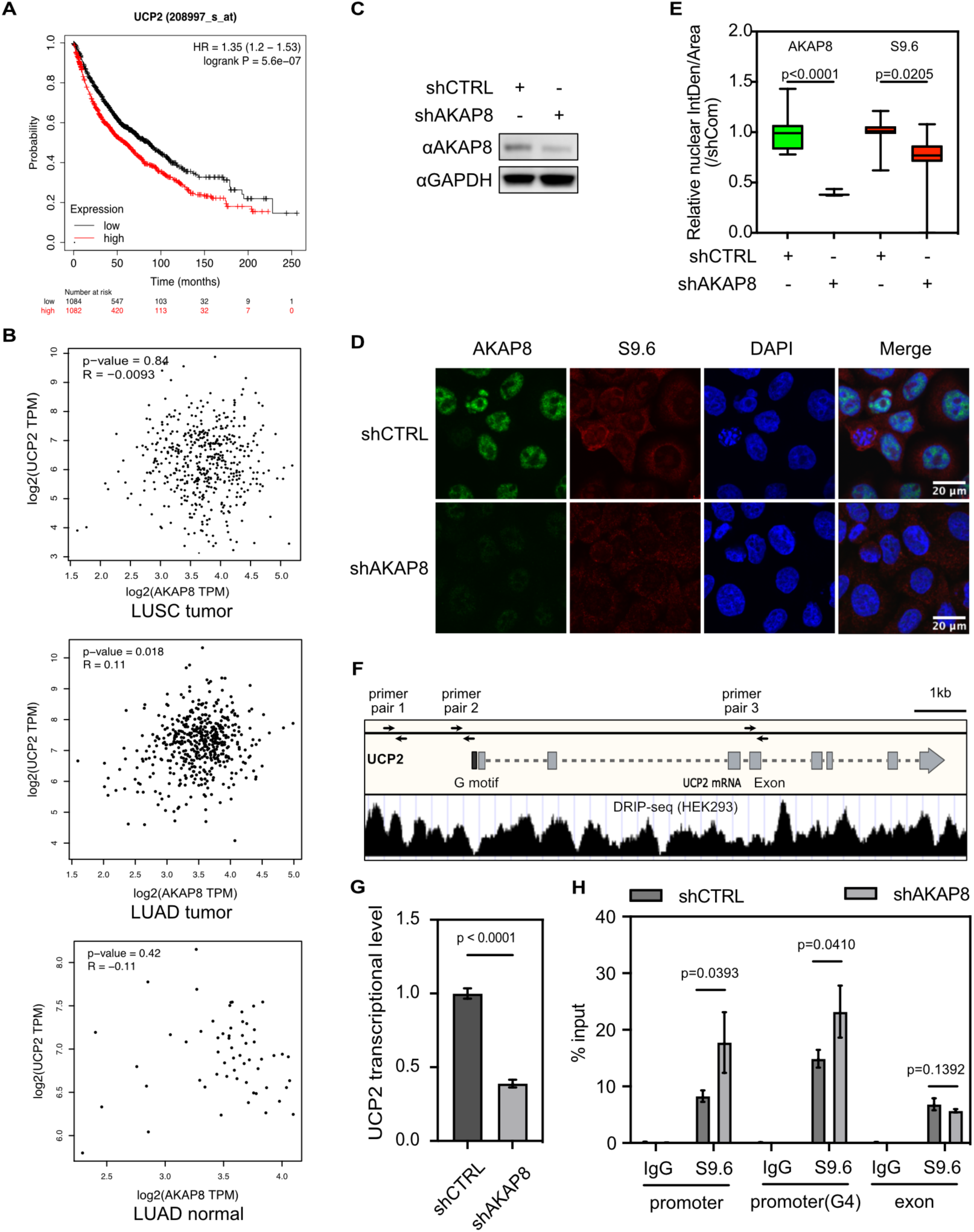
UCP2 regulated by AKAP8 served as contributing factor to lung adenocarcinoma. A, Survival probability with high and low UCP2 expression. B, Correlation analysis of AKAP8 and UCP2 in LUSC tumor, LUAD tumor and LUAD normal tissue. C, AKAP8 expression in shCTRL and shAKAP8 A549 cell line. D, image of immunofluorescence of AKAP8 and R-loop (S9.6) in shCTRL and shAKAP8 A549 cell lines. E, statistics of relative nuclear signal of AKAP8 and R-loop (S9.6). F, diagram for sketching UCP2 gene, mRNA, DRIP-seq signal. G, relative transcriptional level of UCP2 in shCTRL and shAKAP8 A549 cell lines. H, Estimation of R-loop accumulation in promoter and exon region of UCP2.

We estimated R-loop formation after AKAP8 depleting in A549. AKAP8 signal of its knock down cell was significantly decreased in according with expression level and nuclear S9.6 signal also reduced comparing to control cell (Figure 7C-E). We confirmed mRNA level of UCP2 significantly reduced in AKAP8 knock down A549 cell line (Figure 7G). Further the R-loop level of UCP2 gene was estimated. The results showed R-loop significant accumulated in UCP2 promoter but bare changes in exon in AKAP8 knock down cell (Figure 7FH). Together, these results suggested AKAP8 modulated R-loop of UCP2 promoter and mRNA level of UCP2 may play roles in lung adenocarcinoma progress.

## Discussion

In this study, we first claimed that AKAP family member AKAP8 bind with R-loop and associated with cellular R-loop regulation. N terminal of AKAP8 was responsible for its nucleus location and interaction with DDX5, while depleted AKAP8 contributed to increasing of chromatin associated DDX5. The depletion ofAKAP8 also disturbed transcriptional profile in carcinoma cell line by regulating pathways including RNA metabolism and mitochondrial pathway. AKAP8 was predicted as positive regulator for lung carcinoma R-loop plays critical roles in biological processes including transcription initiation/termination, DNA replication, DNA damage/repair. It is also not to be neglected in immunoglobulin gene class switch, genome instability and telomere maintenance (Petermann et al. 2022) (Niehrs and Luke 2020) (García-Muse and Aguilera 2019) (Wigton and Ansel 2021) (Toubiana and Selig 2018). So it is worthy to deeply dig function of R-loop associated proteins. Basing of R-loop antibody S9.6, researches employed pull-down subsequently mass spectrometry technology to identify R-loop proteins in various cells (Cristini et al. 2018) (Wu et al. 2021). Several groups employed inactive RNase H or HBD domain (RNA:DNA hybrid domain from RNase H), both of them capable of binding with RNA:DNA hybrid, combining proximity labeling to explore R-loop interacting proteins. HBD fusion with an engineered variant of APEX2 (ascorbate peroxidase) was employed to develop RNA:DNA proximity proteomics termed as RDProx and identify R-loop interactome (Mosler et al. 2021). Inactive RNase H (D210N, RHΔ) fused an engineered biotin ligase TurboID to identify repertoire of site specific R-loop modulators (Yan et al. 2022). However, S9.6 immunoprecipitation was restricted R-loop interactors identification under in vitro condition since the IP process was conducted by using cell extraction. For RDProx and RHΔ-TurboID method, they were beyond this problem however with their own limitation. Biotin-phenol transferase activity of APEX is dependent on H_2_O_2_ meaning that this system has routine of H_2_O_2_ incubation. This may affected redox balance of cell and triggers metabolism changes leading to R-loop and its interactors disturbance. RNase H1 is monomer in cell which provides advantage for exogenous RHΔ without bother of endogenous RNase H1 for associated protein analysis. However, as a large protein molecular instead of peptide, inactive RNase H1 has its own interactors may cover or shelter the R-loop interactors even though control experiment set.

Taking these into consideration, we employed HBD, for less instinct interactors and TurboID, for harmless proximity label for cell, to establish HBD-TurboID proximity labeling system and to label R-loop interactors. We had verified this system by labeling and detection of previous known R-loop binding protein DDX5 and DHX9. Further we identified new R-loop associated protein AKAP8 by this system (Figure 1, Figure 2). By HBD-TurboID proximity labeling and Sa affinity, we used 20% volume of cell extraction for S9.6 immunoprecipitation as affinity reaction system and achieved more obvious signal of target protein by western bloting in same experimental condition. This method for detection of weak binding R-loop protein appear to be advantageous comparing with traditional S9.6 immunoprecipitation method. Nevertheless, one thing can not be neglected that both S9.6 and HBD showed slight preference for dsRNA, like 1/25 affinity preference of S9.6 binding with RNA:DNA and 1/(16-127) for that of HBD (Bou-Nader et al. 2022) (Nowotny et al. 2008) (Wang et al. 2021). There is possibility that R-loop interactors identified by using two methods may also contain dsRNA binding proteins.

AKAP8 was firstly reported before thirty years but only about sixty researches were indexed. In the beginning the function of AKAP8 was focused on its interacting with cyclins, regulation on mRNA stability and splicing (Arsenijevic et al. 2004) (Jungmann and Kiryukhina 2005) (Kvissel et al. 2007). In recent ten years, AKAP8 was reported to regulation epigenetics modification of H3K4 methylation by functionally association with MLL complexes and gene expression in embryonic stem cells (Jiang et al. 2013) (Bieluszewska et al. 2018). Its roles in signal transduction of ERK, cAMP and DNA damage was also addressed by several research groups (Asirvatham et al. 2021) (Kong et al. 2023) (Chan et al. 2023). It was more attractive that the function of AKAP8 in tumorigenesis associated RNA splicing regulated by its phase-liquid separation. Expression of AKAP8 was reported to be positively associated with growth of breast cancer cell maybe by possibility of regulation of inflammatory response and apoptosis gene expression as well as regulation of splicing for tumorigenesis gene (Li et al. 2020a). Another report showed that AKAP8 antagonized with splicing activity of hnRNPM to prevent EMT (epithelial-mesenchymal transition)-promoting splicing in breast cancer cell (Hu et al. 2020). N terminal IDR (intrinsically disordered region) 101-210aa of AKAP8 especial Tyr site was regarded as the pivot site responsible for phase-liquid dynamic status in nucleus. Mutation of IDR Tyr or deletion of IDR changed phase-liquid balance of nucleus AKAP8 in carcinoma cell resulting in disturb of RNA splicing (Li et al. 2020a). N terminal region 1-100 aa of AKAP8 was reported to interact with RNA processing proteins including splicing factors, RNA helicase and hnRNP by HA IP MS method (Hu et al. 2016). Therefore, N terminal of AKAP8 was regarded as key region for RNA splicing regulation. This region was founded to dominate AKAP8 nucleus distribution as well as interaction with DDX5(Figure 4). However, whether this region is sufficient to regulate biofunction of DDX5 in cell is still needed to be determined.

AKAP8 contains two ZF (zinc finger)domains in its middle region proposed function including mediation of chromatin condensation and RNA motif binding (Collas et al. 1999) (Hu et al. 2016). These motif was supposed to involve in RNA splicing by direct binding with pre-mRNA. We demonstrated that AKAP8 bind with R-loop and regulated its level in carcinoma cell line (Figure 2, Figure 7). It was suggested that complicated regulatory function of AKAP8 in RNA splicing. moreover, R-loop has interference with splicing process. The fact AKAP8 associating with R-loop and regulating its level make it as a more complicated mechanism.

There is less doubt that AKAP8 is closely related to tumorigenesis and progress. Knock down of AKAP8 significantly reduced cell growth and tumorigenicity of breast carcinoma (Li et al. 2020a). AKAP8 was considered as suppressor of tumor metastasis by antagonizing EMT process (Hu et al. 2020). dynamically interacted with Cx43 (connexin 43) and competitively separated from cyclinE1/E2 to regulate G1/S conversion in lung carcinoma cells (Chen et al. 2020) (Chen et al. 2016). Our experiment also revealed that AKAP8 contributed to cell growth of lung carcinoma cell line (Figure). Besides, Transforming growth factor beta(TGF-beta) is considered one of most main driver of fibrosis in progressive interstitial lung disease IPF (Idiopathic pulmonary fibrosis) (Young et al. 2023). RNA-seq results showed that knock down of AKAP8 disturbed TGF-beta/SMAD3 regulating patten (Figure S3C). The regulation of TGF-beta by AKAP8 was also reported in previous research (Li et al. 2020a). Therefore, it is not to be neglected the roles of AKAP8 in lung disease.

In this study, we offered evidence that knock down of AKAP5 increased nucleus DDX5 level. DDX5 as a prognostic marker involves in proliferation and tumorigenesis of NSCLC (non-small cell lung cancer) cells through activating the β-catenin signaling pathway (Wang et al. 2015). Further evidence showed depletion of DDX5 reduced growth and mitochondrial dysfunction in chemoresistant SCLC (small cell lung cancer) cell line by possibility of reduction of intracellular succinate which serves as direct electron donor to mitochondrial (Xing et al. 2020). DDX5 reduced expression in various cancer cell lines and tumor xenografts under hypoxia condition and rescued DDX5 in hypoxia showed R-loop levels accumulation further (Leszczynska et al. 2023). The resolution activation of DDX5 on R-loop in cell was reported only years ago and several co-regulators were also focused in recently as introduced in this text. post translation modification of DDX5 impaired its biofunction in R-loop regulation. RGG/RG motif of DDX5 methylated by protein arginine methyltransferase 5 (PRMT5) was not necessary for R-loop resolution activity but was required for interacting of DDX5 with XRN2 exoribonuclease to resolve R-loop (Mersaoui et al. 2019). DDX5 was phosphorylated at tyrosine residue in cell and was a cellular target of p38 MAP kinase. The phosphorylation of DDX5 showed effects on its RNA unwinding activity (Yang et al. 2005). DDX5 was also physiological substrate of PAK5 (P21-activated kinase 5) phosphorylated DDX5 at Thr69. DDX5-phosphorylation-mediated lysine 53 (K53) SUMOylation enhanced DDX5 stability and both two modification of DDX5 facilitated Drosha/DGCR8/DDX5 complex formation (Li et al. 2021). However there was no direct evidence that phosphorylation of DDX5 involved in R-loop resolution. AKAP8 is kinase anchoring protein and associated with kinase subcellular spatial orientation. Whether AKAP8 interacting with DDX5 direct DDX5 phosphorylation and play roles in regulating cellular R-loop level is an attractive question. It is also interesting that DDX5 interacted with UCP2 in NSCLC cells and affected tolerance of cell to chemotherapy drugs by AKT/mTOR signaling (Yang et al. 2021). In this study, we speculated that UCP2 transcription was down regulated but R-loop level of DD5 gene promoter was up regulated by knock down of AKAP8 in carcinoma cell line (Figure). Therefore, AKAP8, DDX5, R-loop and UCP2 showed a complex regulating net to modeling metabolism of lung carcinoma cell.

## Material and method

### Cell and cell culture

HEK293T human embryonic kidney, A549 human lung adenocarcinoma, HeLa human ovarian carcinoma, U2OS human osteosarcoma cell lines and UMSC umbilical cord mesenchymal stem cell were used in this study. Generally, cell were cultured in DMEM medium supplemented with 10% fetal bovine serum, 1% penicillin/streptomycin in at 37°C in a humidified atmosphere with 5% CO_2_. Cell were cultured to a confluence of about 90% and then passage by 1:3∼5 to fresh complete medium. Cells were regularly confirmed mycoplasma free before experiment.

### Plasmid construction

pLKO.1-TRC-shRNA plasmid were constructed by restriction enzyme digestion and ligation. Oligo of hair pin shRNA of AKAP8 containing flanking Age I and EcoR I enzyme site sequence were synthesized (shCTRL oligo: 5’3’; shAKAP8-CDS oligo: 5’ CCGGGCCAAGATCAACCAGCGTTTGCTCGAGCAAACGCTGGTTGATCTTGGCTTTTTG3’, 5’ AATTCAAAAAGCCAAGATCAACCAGCGTTTGCTCGAGCAAACGCTGGTTGATCTTGGC3’; shAKAP8-3’UTR oligo: 5’CCGGGCTGAAGTACATTGTCCTTAGCTCGAGCTAAGGACAATGTACTTCAGCTTTTTG3’, 5’AATTCAAAAAGCTGAAGTACATTGTCCTTAGCTCGAGCTAAGGACAATGTACTTCAGC3’). The sense and antisense oligo were mixed and heated to 95℃ and annealed to base pair complementary by immediately transfer to ice or programmed gradient cooling to 25℃. pLKO.1-TRC-shRNA empty plasmid were digested by Age I and EcoR I, recycled and ligated with annealed shRNA oligo. pLKO.1-SHC002 (Sigma-Aldrich, #SHC002) was as control shRNA plasmid.

Expression plasmid was generally constructed by seamless method or blunt end ligation. pCR3.1 backbone, AKAP8 with plasmid flanking overlap sequence were amplified and then homologous recombination according to guide of seamless kit instruction to construct pCR3.1-HA-AKAP8. Truncated HA-AKAP8 expression plasmid were constructed by PCR amplified using pCR3.1-HA-AKAP8 as template and blunt end ligation. pCMV3-DDX5-FLAG were bought from Sino Biological company (#HG16175-CF). Truncated DDX5-FLAG expression plasmid was constructed by PCR and blunt end ligation.

pGEX-4T-AKAP8 expressing GST-AKAP8, and pET28a-DDX5 expressing His-DDX5 were also constructed by seamless method. The truncated expression plasmid was constructed by PCR and blunt end ligation.

pLV3-CMV-TurboID-NLS-FLAG were originally bought from MiaoLingBio company. Kozak sequence were added after CMV before TurboID and used to construct HBD-TurboID plasmid. HBD with overlap sequence and pLV3-TurboID-NLS-FLAG backbone plasmid were amplified to homologous recombined as pLV3-HBD-TurboID-NLS-FLAG.

### Lentivirus package and infection

293T cell were passage and cultured over night to confluence of 80% in 6 well plate. The cell transfection were carried out for lentivirus package by using following plasmid: 1 μg pLKO.1-shRNA, 0.75 μg psPAX2, 0.25 μg pMD2.G mixed in 100 μL FBS free DMEM medium. 4 μg PEI in equal volume DMEM was added into plasmid and gently mixed, incubated for 10 min at room temperature. Mixture of plasmid and PEI were dropwise addition to transfect 293T cell. Virus were harvested after 48 h. For infection, target cell were passage and cultured over night to confluence of 80% in 6 well plate. Medium were discarded and 1 mL virus and 1 mL fresh complete medium were added with mixture of final concentration of 8 μg/mL polybrene. Medium was replaced by 2 μg/mL Puro medium after 48 h culture. The selection were lasted at least for 5 days for stable cell line.

### Biotin labelling and Sa affinity precipitation

Biotin stock was prepared at concentration of 100 mM in dimethyl sulfoxide (DMSO). The cell were cultured to confluence about 90%. Biotin stock was diluted by serum medium and directly added to medium at final concentration of 500 μM for 10 min unless indicated otherwise. To terminating the label proceed, cell was transferred to ice, discarding the medium and washed by pre frozen PBS for 3 times. The cell was scraped from plate and harvest by centrifuging under 4°C 500 g for 5 min. Supernatant was removed. The pellet of about 10^7^ cell was resuspended and lysed by 1 mL WCE buffer (50 mM Tris pH 8.0, 150 mM NaCl, 0.5% NP-40, 1 mM EDTA and 10% glycerol). After 15 min rotation in 4°C refrigerator, whole cell lysate (WCL) was clarified by centrifuging 4°C 12000 g for 10 min. The total protein concentration of WCL was estimated by BCA protein assay. WCL containing 0.2 mg protein was incubated with 15 μL Sa magbeads for 1.5 h rotation under 4°C. Beads was subsequently washed twice by WCE buffer, once by 1 M KCl, once by 0.1 M Na_2_CO_3_, once by 2 M urea/10 mM Tirs pH 8.0, recovered by twice WCE buffer washing. For western blotting, the slurry was boiled at 95°C for 5∼10 min by adding 30 μL 1*loading buffer.

### RNA:DNA hybrid (R-loop) associated protein immunoprecipitation

R-loop associated protein of cell was precipitated by S9.6 referring to previous reports. Briefly, 10^7^ cell were harvest and washed twice by pre cold PBS. 1 mL WCL buffer was added and fully suspended by vortex following 30 min incubation on ice. After high speed centrifugation, supernatant were collected for IP. Antibody was incubated for 2 h with 15 μL pre balanced proteinA/G beads in 100 μL WCL buffer per IP reaction. 500 μL WCL was added into beads and incubated for 2 h in 4°C condition with gently upside down mix. Beads were separated and supernatant was discarded. Beads was washed by 500 μL WCL for 3 min at least 6 times. The beads slurry was resuspended by 30 μL SDS-loading buffer and boiled under 95°C for 10 min. the IP products was separated by SDS-PAGE and target protein was estimated by western blotting. For RNase H or RNase A test group, 5U RNase H or 10 μg RNase A were added per IP reaction and incubated at 37℃ overnight.

### Immunofluorescence

Cell was cultured into confluence about 90% and enzymatic digested by trypsin, seeded by 1:5 in 6 or 12 well plate before 24h of immunofluorescence. The 20 mm slide was pre-autoclaved and placed in well before seeding cell. For transfecting, cell was seed by 1:8 and cultured for 24 h. Transfecting of cells was conducted according to method mentioned in this section. After 36 h culture, the following operation was carried out. The plate was washed by PBS, fixed by 4% PFA for 15 min, stop fixation process by 2mg/mL Glycine for 10 min twice, permeated by 0.2% Triton X-100 for 10 min. Following PBS washing twice, cells were blocked by 3% BSA/PBS for 1h at room temperature. After PBS washing thrice, cells were incubated with primary antibody diluted by 3% BSA/PBS for 1 h at room temperature. After washing thrice by wash buffer (1% BSA, 0.05% Tween 20, PBS) thrice, cells were incubated with fluorescence labeled second antibody for 1 h. After washing thrice by wash buffer, cells were incubated with DAPI for DNA stain. The slide were sealed for imaging by Nikon A1. For R-loop immunofluorescence, cell were fixed and permeated by pre-cold methanol for 10 min at -20℃ and acetone for 1 min at room temperature. After washing thrice by PBS, the cells were blocked and incubated with antibody.

### Antibody used in this study

S9.6 (Merck, #MABE1095), anti-AKAP8 (Zenbio, #R26398), anti-AKAP8L (Zenbio, #160665), anti-DDX5 (Cell signaling technology, #9877; Proteintech, #67025-1-Ig), anti-DHX9 (Zenbio, #382331), anti-H2A.X (Zenbio, #201082-7G9), anti-FLAG (Sigma-Aldrich, #F1804; Proteintech, #80010-1-RR), anti-HA (Proteintech, # 66006-2-Ig), anti-GAPDH (Proteintech, #60004-1-Ig), anti-H3 (Proteintech, #68345-1-Ig), Normal Rabbit IgG (Cell signaling technology, #2729), Mouse IgG (Proteintech, # B900620), HRP-Goat antibody (Proteintech, SA00001-2, SA00001-3), fluorescence crosslinked donkey antibody (Proteintech, # SA00013-5, SA00013-6, SA00013-7, SA00013-8).

### DRIP to enrich R-loop DNA

Cells were cultivated into confluence of 90% and medium were discarded. Cells were washed by PBS twice and scraped from dish placing on ice. After centrifugation at 4℃ 500g for 5 min, 10 cm dish cultivated or 10^7^ cell were harvested and resuspended by 0.5 mL lysis buffer (10mM Tris pH 8.0, 1% SDS, 2mM EDTA, 100mM NaCl) with 50 μg/mL Protein K. after incubation at 37℃ overnight, final concentration of 20 μg/mL RNase A (DNase free) were added and incubated for 30 min. Add one volume phenol/chloroform extract liquid, vertex and rest for 10 min at room temperature. Transfer to phase gel lock tube and centrifugate at 12000 rpm for 5 min, transfer liquid to new tube. Add 1/10 volume 3 M NaAc (pH 5.2) and 2 volume alcohol. After DNA depositing at -20℃ for at least 30 min, centrifugate at 12000 rpm for 10 min and discard supernatant. DNA was resuspended by 1 mL 80% alcohol and then centrifugated at 10000 rpm for 10 min. supernatant was aspirated totally and DNA was air-dried. About 1 mL TE buffer (10 mM Tris–HCl pH 8.0, 1 mM EDTA) was added and DNA was totally resolved for DNA concentration measurement. Diluted DNA concentration to 0.5 mg/mL. DNA was fragmented into 200-1000 bp by ultrasonication. 300 μL DNA (150 μg) was placed in sonication tube (Diagenode, Bioruptor Pico). The parameter of sonication instrument was set at 4℃ 15 s/45 s (work/off) for 8 cycles. 50 μg DNA fragment was diluted by IP buffer (10 mM Na_2_HPO_4_ pH 7.0, 140 mM NaCl, 0.05 % Triton X-100) to 500 μL was used for DRIP. 15 μL Protein A/G beads used for one DRIP sample. Protein A/G beads was pre-blocked by 0.5% BSA/PBS for 2 h and washed by IP buffer for twice. Protein A/G beads was incubated with 2 ug S9.6 in 100 mL IP buffer at 4℃ for 2 h by upside down mix. 50 μg DNA fragment was diluted by IP buffer up to 500 mL. S9.6 protein A/G beads suspend slurry was added into DNA fragment. Before this, 5 μL DNA was aspirated as input. The IP component was mixed by gently upside down at 4℃ for 2 h. beads slurry was separated and supernatant was aspirated. Beads slurry was washed with 1 mL IP buffer over five times. the beads slurry was resuspended by 200 μL PK buffer (50 mM Tris–HCl pH 8.0, 10 mM EDTA, 0.5 % SDS) with 1 μL 20 mg/mL protein K and incubated at 55℃ for 4 h. IP DNA and input diluted by IP to total volume of 200 μL was purified by phenol/chloroform protocol mentioned in this method. Purified DNA was resolved by 50 μL TE buffer for further experiment.

### Full length transcriptome

Cells were cultured to influence of 90% in 6 well plate. Medium was discarded and cells were washed by PBS twice. 1 mL TRIZOL was added for lysis of cell. The lysate was transferred to tube and incubated at room temperature for 5 min. 0.2 mL chloroform was added and drastic mixed and then placed for 10 min. One volume of isopropanol was added and mixed upside down. after centrifugation of 12000 rpm for 10 min, two volume alcohol was added to supernatant. After DNA depositing at -20℃ for at least 30 min, centrifugate at 12000 rpm for 10 min and discard supernatant. DNA was resuspended by 1 mL 80% alcohol and then centrifugated at 10000 rpm for 10 min. Supernatant was aspirated totally and DNA was air-dried. 50 μL distill water was added to resuspend total RNA at 65℃ for 10 min. Oxford Nanopore Technologies(ONT) full length RNA-seq and analysis was carried out by Biomarker Technologies Co., LTD.

### qPCR

cDNA was synthesized by reverse transcription using of 1μg total RNA as template according to manufacture instrument of HiScript III RT SuperMix for qPCR (Vazyme, #R323-01). cDNA was generally diluted 10 folds by distill water. For IP DNA qPCR, DNA was resuspended in 50 μL distill water. qPCR was carried out using of PCR mix, DNA template, primer according manufacture instruments of TB Green Fast qPCR Mix (Takara, #RR430) and CFX96 Real-Time System (BIO-RAD, C1000 Touch). The Ct value was used for relative abundance calculation by 2^(-ΔΔCt)^ method.

### Cell colony formation

Cells were cultured in complete medium to logarithmic phase and enzymic digested from dish. Cells were then inoculated in 6 well plate by 1000 cell per well. After 24 h, plasmids were transfected by PEIpro and medium were refreshed after 24 h. cell were continue cultured for another 10 days. Medium were discarded and cell were washed by PBS twice. Cells were fixed by adding 1 mL methanol for 30 min. methanol was discarded and cell were stained by 0.1% crystal violet for 3 min. Plate were washed by PBS to clear for cell colony imaging.

### Isolation of cytoplasm, nucleoplasm, chromatin associated fraction

10^7^ cultured in 10 cm dish was washed by pre-cold PBS twice. Cells were scraped and harvested by centrifugation at 4℃ 500g for 5min. Cells were resuspended by 0.5 mL cytoplasmic lysis buffer (50 mM Tris-HCl pH 8.0, 140 mM NaCl, 1.5 mM MgCl_2_, 0.5% NP-40, 1 mM DTT and protease inhibitor) and lysed for 5 min in ice. After centrifugation of 800 g for 2 min at 4℃, supernatant was harvested as cytoplasmic fraction after high speed centrifugation. The pellet was washed by cytoplasmic lysis buffer without NP-40, centrifuged and then resuspended in 50 μL nucleoplasmic lysis buffer 1 (20 mM Tris-HCl pH 7.9, 75 mM NaCl, 0.5 mM EDTA and 50% (v/v) glycerol) totally. 0.5 mL pre-cold nucleoplasmic lysis buffer 2 (20 mM HEPES-KOH pH 7.6, 300 mM NaCl, 0.2 mM EDTA, 7.5 mM MgCl_2_, 1% (v/v) NP-40, 1 M urea and protease inhibitor) was added to suspension, vortexed and incubated for 15 min on ice. The supernatant was gently harvested by centrifugation at 4℃ 2000 g for 4 min as nucleoplasmic fraction after high speed centrifugation. The pellet was quickly washed by nucleoplasmic lysis buffer 2 and harvested by centrifugation of 13000 g at 4℃ for 4 min. the supernatant was discarded and pellet was resuspended by 0.5 mL high salt buffer (50 mM Tis pH 8.0, 500 mM NaCl and protease inhibitor) with 250 U Turbo DNase and incubated for 30 min at 37℃ as chromatin associated fraction.

### Bioinformatic analysis and statistics analysis

Statistic expression of AKAP8 and graph plot in pan-cancer/normal tissue and different types of lung cancer/normal tissue were analyzed by UALCAN platform. Correlation between gene and survival in lung cancer were analyzed by Kaplan-Meier Plotter with default parameters. Dimensionality reduction of DEG set from RNA-seq, gene correlation, and graphic plotting were analyzed by GEPIA2 platform. T-test and p value were estimated by Graphpad Prism 9.

## Compliance and ethics

The author(s) declare that they have no conflict of interest.

## Acknowledgements

This work was supported by the Guangzhou National Laboratory Research Fundation (YW-YWYM0401) and National Natural Science Foundation of China (31900655). It was grateful to Feng Zhou (Guangzhou Laboratory) for regent ordering, Ms. Lanlan Zhang (Guangzhou Laboratory) for help of confocal microscope operation, Dr. Yaping Chen (Chinese Academy of Science) for kind gift of pCR3.1-HA-AKAP8 plasmid, Ms. Di Wu (Guangzhou Medical University) for kind gift of UMSC cell extraction, Ms. Lin Lu and Prof. Yan Wang (Sun Yat-sen University) for funding (31900655) managements and Prof. Dajiang Qin (The Fifth Affiliated Hospital of Guangzhou Medical University) for helpful advisement.

## Figure and table legends

**Figure S1. AKAP8 bind with R-loop.** A, AKAP8 interactome contains RNA process (hnRNPs, Exoscs), chromatin associated (HIST, DPY30) and R-loop associated proteins (RPA, DDX). B, S9.6 IP for estimation of AKAP8 and AKAP8l bind with R-loop. Input: 293T whole cell extraction.

**Figure S2. AKAP8 bind with DDX5.** A, Co-IP analysis interaction of endogenous expressed AKAP8 and DDX5. B, Co-IP analysis interaction of half-exogenous expressed AKAP8 and DDX5. C, interaction estimation of AKAP8 and DDX5 after benzonase treatment. D, interaction estimation of exogenous AKAP8 and DDX5.

**Figure S3. AKAP8 regulated cell metabolism.** A, PC analysis of shCTRL and shAKAP8 cell lines. B, Alternative polyadenylation (APA) site motif clustering. C, clustered heatmap of significant TF’ in shCTRL and shAKAP8 samples. D, GSEA analysis revealed AKAP8 knock down regulated RNA degradation.

**Table S1. Plasmid used in this study.**

**Table S2. Primers for qPCR used in this study.**

