## Supplemental File for "AKAP8 interacting with DDX5 to regulate R-loop balance involves in lung carcinoma cell growth"

Figure S1

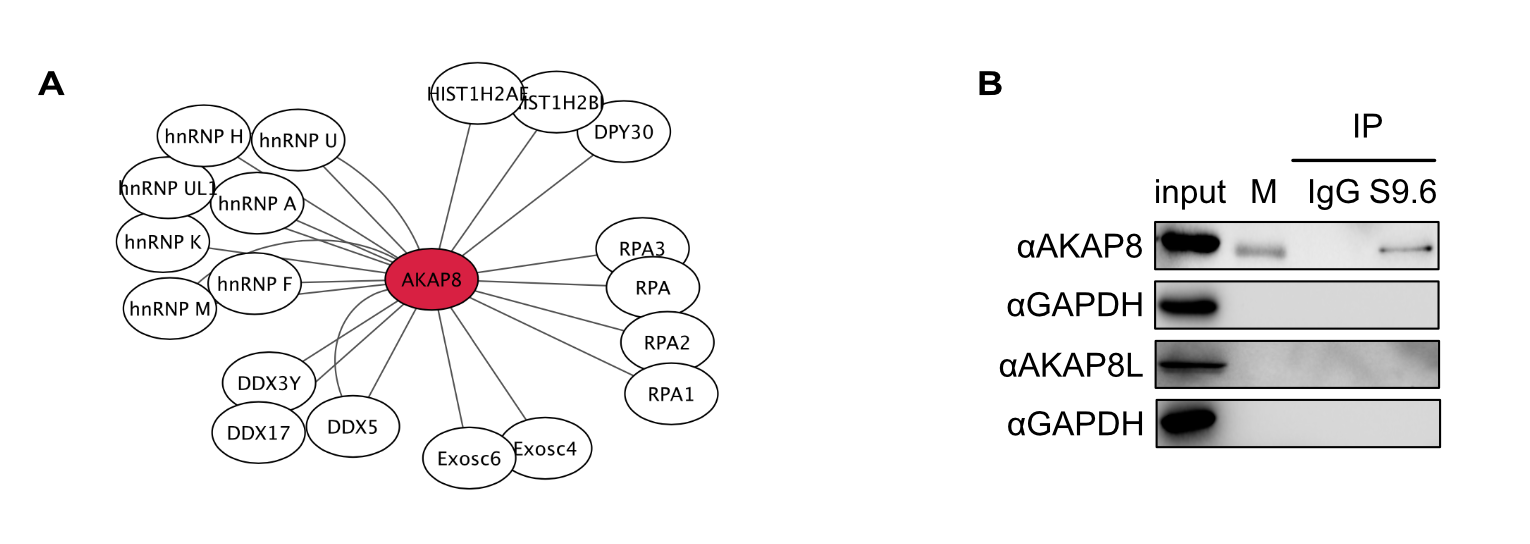

Figure S2

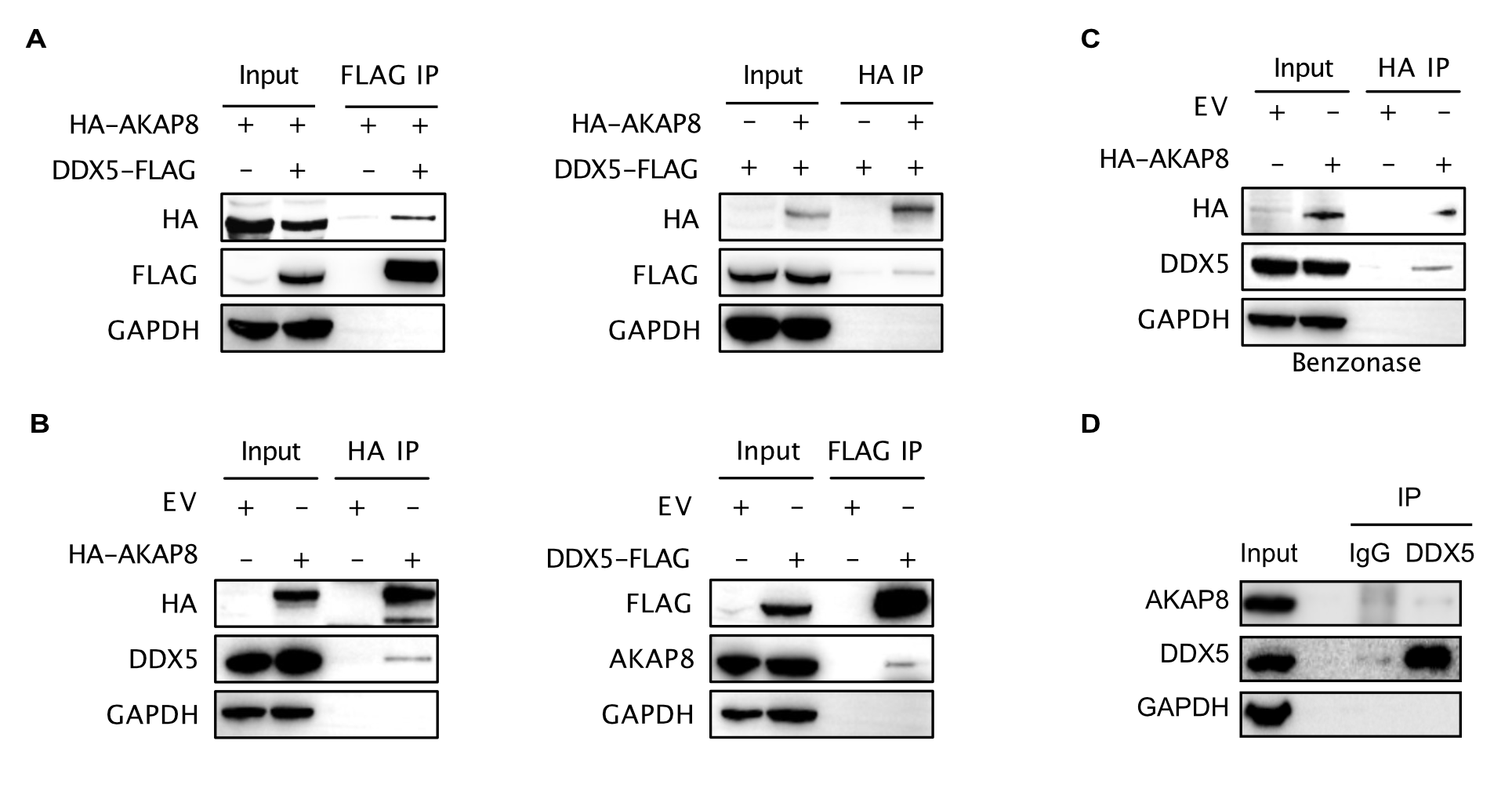

Figure S3

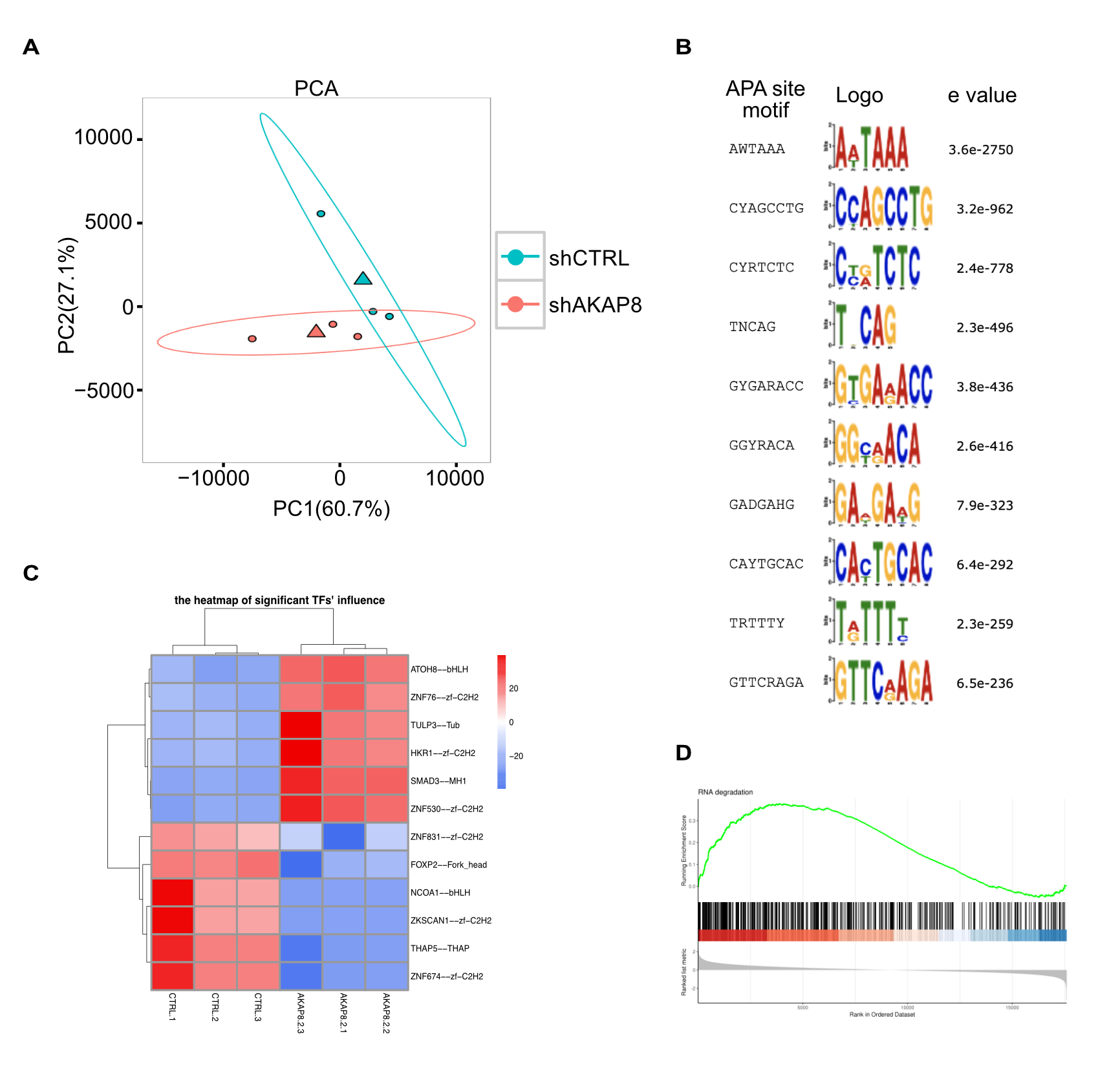

Table S1

**Plasmid used in this study**

pLKO.1-shCTRL

pLKO.1-shAKAP8-CDS

pLKO.1-shAKAP8-3UTR

pCR3.1-HA-EV

pCR3.1-HA-AKAP8

pCR3.1-HA-AKAP8 Δ1

pCR3.1-HA-AKAP8 Δ2

pCR3.1-HA-AKAP8 Δ3

pCR3.1-HA-AKAP8 Δ4

pCMV3-FLAG-EV

pCMV3-DDX5-FLAG

pCMV3-DDX5-FLAG Δ1

pCMV3-DDX5-FLAG Δ2

pCMV3-DDX5-FLAG Δ3

pCMV3-DDX5-FLAG Δ4

pGEX-4T-1

pGEX-4T-AKAP8

pET28a-His-DDX5

pLV3-CMV-TurboID-NLS-FLAG

pLV3-EV-NLS-FLAG

pLV3-TurboID-NLS-FLAG

pLV3-HBD-TurboID-NLS-FLAG

Table S2

| **Primer for qPCR used in this study** | | | |
| --- | --- | --- | --- |
|  | Gene | Sequence | |
| cDNA qPCR | AKAP8 | Forward | GAAGCAGTTCCAACTTTACGAGG |
|  |  | Reverse | CAGAGTTCATCCTCACCCTTG |
|  | GAPDH | Forward | AAGGTGAAGGTCGGAGTCAA |
|  |  | Reverse | AATGAAGGGGTCATTGATGG |
|  | UCP2 | Forward | TGGTCGGAGATACCAAAGCACC |
|  |  | Reverse | GCTCAGCACAGTTG ACAATGGC |
|  | DDX5 | Forward | GCCGGGACCGAGGGTTTGGT |
|  |  | Reverse | CTTGTGCTGTGCGCCTAGCCA |
|  | TMX3 | Forward | ATCTGGGGCTCTAATTCGGC |
|  |  | Reverse | AGTGTCACATACTCAGGAACCA |
|  | DERA | Forward | CAAGGCTGCAGGCTGT AATA |
|  |  | Reverse | CATCTTCCACAGCCAATCTG |
|  | DYNC1LI2 | Forward | CTCCATTCTGAGCGAAGTGTC |
|  |  | Reverse | GGCCTTTGTGGTACAAGTCTC |
|  | SEC11A | Forward | TGTCTCATCGGCACTAATGATCT |
|  |  | Reverse | AGCACCACTACAATCGGACTT |
|  | VPS4B | Forward | ATGTCATCCACTTCGCCCAAC |
|  |  | Reverse | TTGCTTGGCTTTATCACCCTG |
|  | MYBPC1 | Forward | GACTGGACCCTTGTCGAAACT |
|  |  | Reverse | TCTTCACCAACTTTCACTGTTCC |
|  | NDUFA4L2 | Forward | ATGATCGGCTTAATCTGCCTG |
|  |  | Reverse | TCCGGGTTGTTCTTTCTGTCC |
| ChIP qPCR | MYC promoter | Forward | TCTTCGGACCTTCTGCAGCCAAC |
|  |  | Reverse | CAGGAGCGTCCGAGGTGCAAG |
|  | MYC promoter (G4) | Forward | CTACGGAGGAGCAGCAGAGAAAGG |
|  |  | Reverse | GTGGGGAGGGTGGGGAAGGT |
|  | MYC exon | Forward | TATGTGGAGCGGCTTCTCGG |
|  |  | Reverse | AAGACCACCGAGGGGTCGAT |
|  | UCP2 promoter | Forward | AGGACTTCATGCTGCGTCCTG |
|  |  | Reverse | TTCCGGAGAGGCTCGGCAAA |
|  | UCP2 promoter (G4) | Forward | TGCACTTAAGACACGGCCCC |
|  |  | Reverse | CACTCCCGTTTCCGTGCGTC |
|  | UCP2 exon | Forward | TGACCATGGTGCGTACTGAGG |
|  |  | Reverse | TCATACAGGCCGATGCGGACA |
